# Accelerated trans-sulfuration metabolically defines a discrete subclass of ALS patients

**DOI:** 10.1101/609925

**Authors:** Qiuying Chen, Davinder Sandhu, Csaba Konrad, Dipa Roychoudhury, Benjamin I. Schwartz, Roger R. Cheng, Kirsten Bredvik, Hibiki Kawamata, Elizabeth L. Calder, Lorenz Studer, Steven. M. Fischer, Giovanni Manfredi, Steven. S. Gross

## Abstract

Amyotrophic lateral sclerosis (ALS) is a disease characterized by progressive paralysis and death. Most ALS cases are sporadic (sALS) and patient heterogeneity poses a formidable challenge for the development of viable biomarkers and effective therapies. Applying untargeted metabolite profiling on 77 sALS patient-derived primary dermal fibroblast lines and 45 sex/age matched controls, we found that ∼25% of cell lines (termed sALS-1) are characterized by upregulated trans-sulfuration, where methionine-derived homocysteine is channeled into cysteine and glutathione synthesis. sALS-1 fibroblasts exhibit a growth defect when grown under oxidative conditions, that can be fully-rescued by N-acetylcysteine. [U-^13^C]-glucose tracing shows that activation of the trans-sulfuration pathway is associated with accelerated glucose flux into the TCA cycle. Based on four metabolites, we developed a support vector machine model capable of distinguishing sALS-1 with 97.5% accuracy. Importantly, plasma metabolite profiling identifies a systemic perturbation of cysteine metabolism as a hallmark of sALS-1. These results indicate that sALS patients can be stratified into distinct metabotypes, differently sensitive to metabolic stress, and provides new insights into metabolic biomarkers for personalized sALS therapy.

## Introduction

Amyotrophic lateral sclerosis (ALS) is a fatal neurodegenerative disease characterized by progressive death of upper and lower motor neurons; symptoms include muscle weakness and atrophy, commonly leading to fatal paralysis within 5 years of disease onset^1^. Approximately 90% of ALS patients have no familial history (a condition termed sporadic ALS; sALS), while the remaining 10% are due to recognized inherited gene mutations^2^. Unfortunately, mechanisms leading to motor neuronal death in sALS are completely unknown and, as a consequence, there are neither viable disease biomarkers nor effective therapies. Past ALS clinical trials have been largely unsuccessful^3^ and currently there are only two approved drugs, Riluzole and Edaravone, both of which only prolong the lifetime of ALS cases by a few months. Conceivably, sALS can originate from multiple mechanistic triggers, and disease stratification (i.e., recognition of distinct disease-driver subtypes) holds potential to uncover targeted new therapies.

As heterogeneity of the sALS patient population presents a major stumbling stone to rigorous mechanistic studies, stratifying sALS has become an area of considerable interest^4, 5^. The worldwide *ALS Stratification Prize - Using the Power of Big Data and Crowdsourcing for Catalyzing Breakthroughs in ALS* was initiated in 2015, aiming to address the problem of ALS patient heterogeneity with consideration of key clinical distinctions, such as rankings on the ALS functional rating scale^6^. Stratification efforts have identified several potential non-standard predictors of disease progression, including plasma uric acid, creatinine and surprisingly, blood pressure that may be related to sALS pathobiology^6^. While past attempts at sALS case stratification has primarily relied on clinical parameters to anticipate disease progression, recognition of disease-associated metabolic types (i.e., *metabotypes*) is anticipated to be more telling, with potential to identify distinct molecular disease mechanisms that inform rationally-targeted therapies.

Altered energy metabolism often precedes the clinical onset of ALS^7^. Moreover, alterations in energy metabolism have been inferred as a potential pathogenic mechanism underlying sALS^8, 9^. It has recently been reported that hypermetabolism in ALS is associated with greater functional decline and accelerated mortality^10^. Notably, as the final products of genes, transcripts, and proteins, metabolites offer the most proximal insights into sALS disease mechanisms. Whereas specific metabolic perturbations that underlie bioenergetic alterations in sALS have remained elusive, understanding this phenomenon will likely inform on disease mechanisms and prove transformational for clinical care.

We sought to investigate whether the metabotype of dermal fibroblasts from sALS patients can enable patient subtyping and disease stratification that may shed light on underlying disease mechanisms. Notably, fibroblasts carry the same genetic composition as neurons, but unlike neural tissue, fibroblasts are readily accessible from patients for *in vitro* culture and analysis. Indeed, prior studies of fibroblast metabolism have revealed bioenergetic alterations in sALS cases^5, 9, 11^. Here, for the first time, we performed untargeted metabolite profiling on fibroblasts from sALS patients and healthy control subjects, in combination with assessments of cell viability and determination of plasma metabolites and gene expression profiles. Studies reveal a unique subset of sALS patients that display a distinct metabotype, typified by accelerated trans-sulfuration pathway-derived cysteine for support of GSH biosynthesis, along with hypermetabolism of glucose. Functionally, cells with this metabotype exhibit increased susceptibility to metabolic stress, which can be prevented by supplementation with the antioxidant N-acetylcysteine. The finding that sALS patients can be classified based on metabotypes associated with cell viability and response to antioxidants offers a potentially critical first-step towards targeted personalized medicines and effective therapy for this newly identified sALS patient metabotype.

## Results

### Untargeted metabolite profiling identifies a distinct sALS metabotype

Table 1 summarizes the clinical characteristics of 77 sALS cases and 45 healthy controls studied herein. Dermal fibroblast cell lines were established from all cases and 1172 quality control-normalized metabolite features were detected in greater than 70% of the samples, in either sALS or control fibroblast groups. Metabolite profiling and non-parametric Mann-Whitney statistical analysis (P<0.05) revealed that 507 of these 1172 recognized features are differentially expressed in sALS fibroblasts. Unsupervised machine learning principal component analysis (PCA) did not completely separate sALS as a whole from the control group (Fig 1B), but a distinct subgroup of 18/77 sALS patient-derived cell lines clustered together, separating from controls based on principal component 1 (PC1), which accounted for 33.9% of total variance.

**Figure 1.**
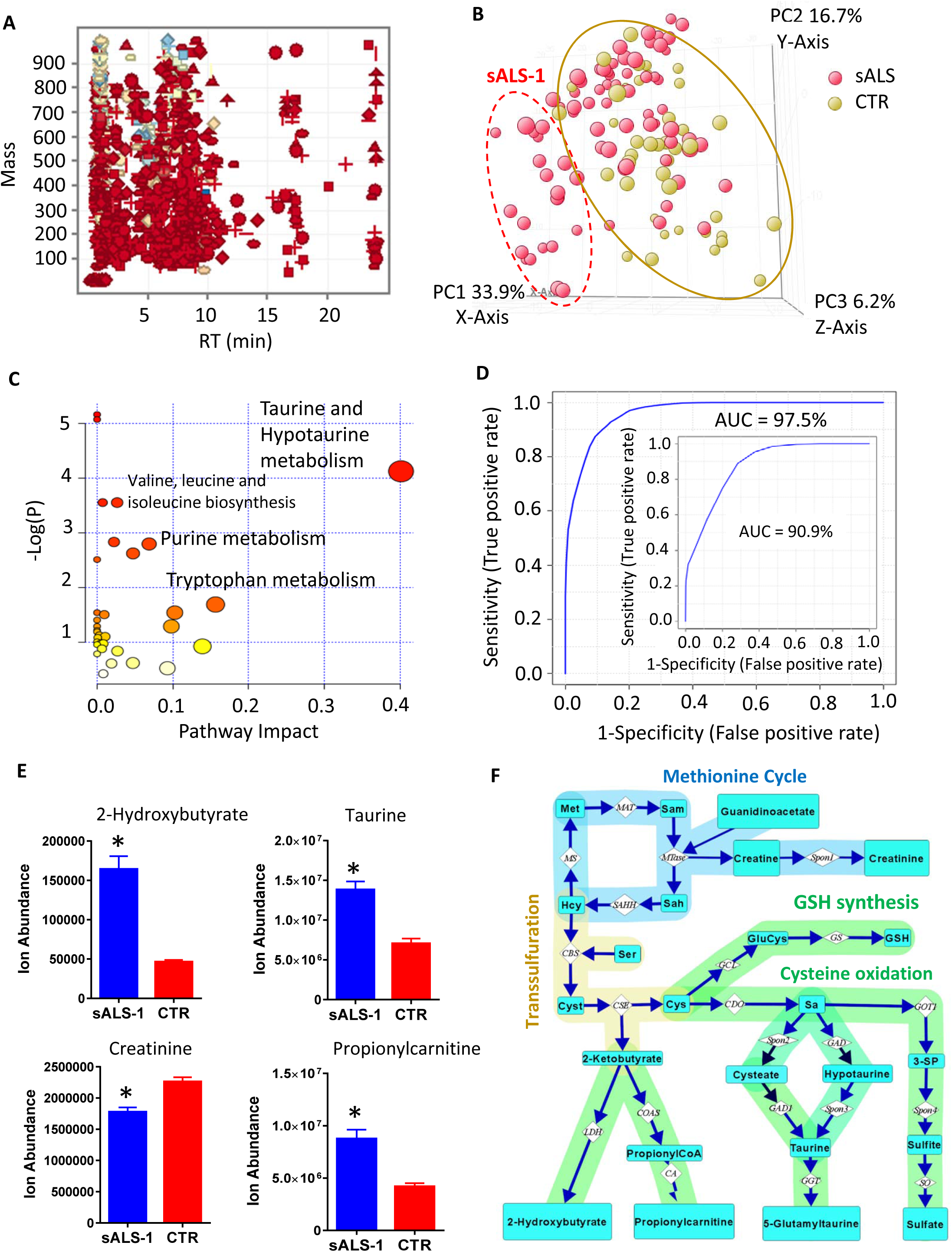
Untargeted metabolite profiling of case-derived fibroblast lines identifies an sALS subgroup with enhanced trans-sulfuration pathway and glucose hypermetabolism. Panel A: mass and retention time (RT) scatter plot of 1165 features detected in greater than 70% of cell cultures in either 77 sALS or 45 control fibroblast groups. Panel B; PCA score plot showing incomplete separation of 77 sALS (red circles) from 45 controls (yellow circles), but a subgroup of 18 sALS patients (identified in the dashed red oval as sALS-1) was readily separated from all controls. Panel C: integrated enrichment and pathway topology analysis of differential metabolites in sALS-1 by Metaboanalyst 3.0, recognizing taurine and hypotaurine in the trans-sulfuration pathway as the most significant pathway impact and lowest matching P-value. Panel D: *ROC* curve derived from SVM model predicting 18 sALS-1 subgroup from a group of 45 randomly selected control fibroblasts. The AUC (i.e. the overall accuracy, specificity and sensitivity) of the prediction is 97.5%. The inset showed the *ROC* curve and AUC for a repeat experiment comprising the same 18 sALS-1 subset vs. 18 randomly selected control fibroblast cell lines. Panel E: Ion abundance of 4 sALS-1 predictor metabolites (2-hydroxybutyrate, taurine, propionylcarnitine and creatinine) from 18 sALS-1 and 45 controls; *, P<0.05 (2-tailed Student’s t-test). Panel F: Schematic representation of metabolites and enzymes involved in the methionine cycle, methylation and trans-sulfuration pathways. Abbreviation of metabolite names and enzymes are as follows: Met, methionine; Hcy: homocysteine; Sam: S-adenosylmethionine; Sah: S-adenosylhomocysteine; Cyst: cystathionine; Sa: sulfoalanine; 3-SP: 3-sulfopyruvate. The enzyme name abbreviations were adapted from the *Kyoto Encyclopedia of Genes and Genomes* (KEGG) pathway. MAT: methionine adenosyltransferase; MS: methionine synthase; SAHH: S-adenosylhomocysteine hydrolase; MTase: methyltransferase; CBS: cystathionine β synthase; CSE: cystathionine γ lyase; GCL: glutamate cysteine ligase; GS: GSH synthase; CDO: cysteine dioxygenase; LDH: lactate dehydrogenase; COAS: propionylCoA synthetase; CA: carnitine acyltransferase; GAD1: glutamate decarboxylase 1; GGT: Gamma-glutamyltransferase; GOT1: glutamic-oxaloacetic transaminase 1; SO: sulfite oxidase; Spon1, Spon2 and Spon3 refer to non-enzymatic spontaneous degradation process.

**Table 1.** Clinical characteristics of the sALS and control cohorts.

| **Subject** | **sALS-Male** | **sALS-Female** | **sALS-Total** | **CTR-Male** | **CTR-Female** | **CTR-Total** |
| --- | --- | --- | --- | --- | --- | --- |
| **N** | 43 | 34 | 77 | 23 | 22 | 45 |
| **BMI** | 26.54 ± 4.55 | 26.04 ± 6.41 | 26.32 ± 5.41 | NA | NA | NA |
| **Age at Symptom Onset** | 56.28 ± 9.92 | 62.44 ± 8.45 | 59.00 ± 9.74 | NA | NA | NA |
| **Age at Time of Skin BX** | 57.65 ± 9.76 | 63.858 ± 0.44 | 60.39 ± 9.66 | 64.83 ± 8.95 | 57.32 ± 5.40 | 61.15 ± 8.27 |
| **\* ALSFRS-R Total at Biopsy** | 32.81 ± 8.61 | 32.38 ± 7.80 | 32.62 ± 8.21 | NA | NA | NA |
| **% Rate of Decline** | 1.20 ± 0.84 | 1.08 ± 0.56 | 1.141 ± 0.72 | NA | NA | NA |
| **#FVC (%)** | 74.78 ± 25.61 | 75.84 ± 29.05 | 75.25 ± 27.00 | NA | NA | NA |

Specified values are means ± S.D.
\*ALSFRS-R = ALS functional rating scale–revised.

To confirm that the sALS subgroup (denoted sALS-1) could be reproducibly distinguished from the control group, we repeated the metabolic profiling in an independent experiment, comparing the same 18 sALS-1 cases vs. a group of 18 randomly selected control lines. Results confirmed 42 metabolites to be differentially-expressed in both the original study and the validation study (Fig S1). Among these metabolites, significant increases were observed for 2-hydroxybutyrate, taurine, and C3-C5 carnitines, along with significant decreases in creatinine and several amino acids. Integrated enrichment and pathway topology analysis of these differentially-expressed metabolites (applying Metaboanalyst 3.0) revealed the taurine/hypotaurine pathway to be the most significantly impacted (Fig 1C).

Using a support vector machine (SVM) learning algorithm and ROC-curve based biomarker analysis, we generated a 4-metabolite linear SVM model (comprising 2-hydroxybutyrate, taurine, propionylcarnitine and creatinine) that recognized the sALS-1 subgroup with an overall accuracy of 97% in the original group (with 1/45 false-positives and 1/18 false-negatives; Fig 1D) and 90.9% accuracy in a repeat analysis. An independent targeted quantitative analysis confirmed the observed decreased level of creatinine and increased levels of 2-hydroxybutyrate, taurine, and propionylcarnitine (Fig 1E).

Remarkably, this group of 4 metabolites share interconnectivity in the methionine cycle, methylation and trans-sulfuration pathways (Fig 1F). Under metabolic stress, supplies of L-cysteine for glutathione synthesis can become limiting, causing the diversion of homocysteine, which normally regenerates methionine, into the trans-sulfuration pathway for production of cystathionine as the first step in cysteine biosynthesis. In this pathway, cystathionine is cleaved to cysteine and 2-ketobutyrate, with the latter species undergoing enzymatic reduction to 2-hydroxybutyrate. An additional fate for 2-ketobutyrate is entry into TCA cycle via addition to propionylcarnitine for fatty acid β-oxidation in mitochondria (Fig 1F). Creatine is synthesized from guanidinoacetate in a S-adenosylmethionine (SAM) dependent reaction^12^ and can spontaneously degrade to creatinine (Fig 1F). Collectively, these untargeted metabolite profiling findings indicate selective trans-sulfuration pathway perturbations in the sALS-1 subset of ALS cases.

### Serine incorporation into *glutathione* (GSH) is increased in sALS-1

We next asked whether the observed increases in trans-sulfuration may be linked to increased glutathione (GSH) synthesis in sALS-1. To test this possibility, sALS-1 and control fibroblasts were grown in 2 mM [2,3,3-^2^H]-serine supplemented medium, followed by a determination of the extent of deuterium enrichment in GSH. This stable isotope tracing experiment studied three sALS fibroblast lines with the highest levels of intracellular 2-hydroxybutyrate, compared with three randomly selected control lines. Notably, serine metabolism to glycine would predictably result in a singly-deuterated glycine molecule for incorporation into *de novo* synthesized GSH. Indeed, sALS-1 fibroblasts in medium supplemented with [2,3,3-^2^H]-serine demonstrated a modest, but significant increase in isotopic incorporation into GSH relative to controls after growth for 1h, 5h, and 24h, (Fig 2A, B). Similarly, glutathione disulfide (GSSG) in sALS-1 fibroblasts showed a significantly elevated serine-derived glycine incorporation, and significantly increase absolute levels of GSSG relative to control fibroblasts after growth for 5h and 24h (Fig 2C, D).

**Figure 2.**
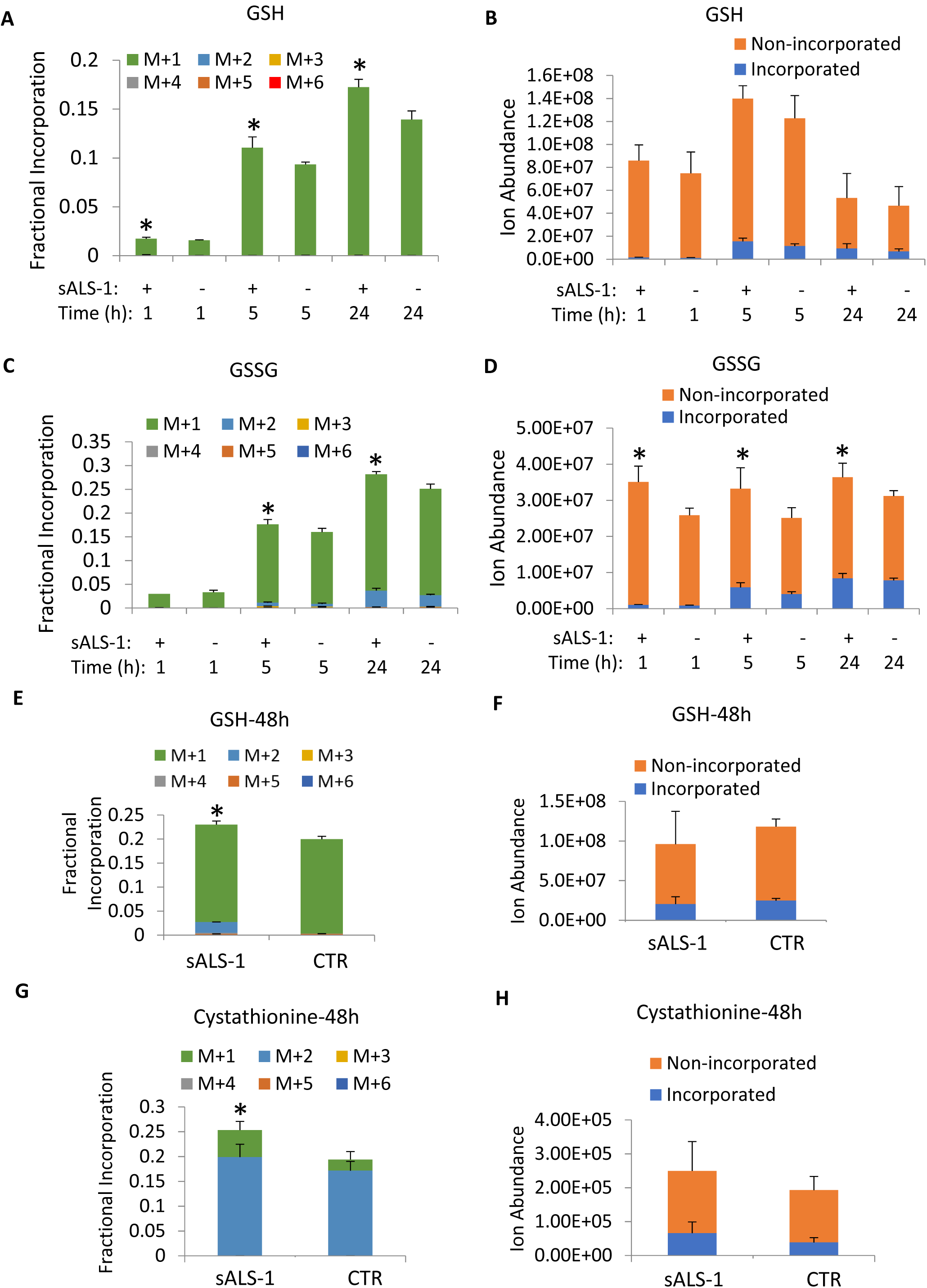
[2,3,3-^2^H]-serine incorporation into GSH, GSSG and cystathionine synthesis is increased in sALS-1. Panel A and B: fractional isotopic enrichment (A) and total ion abundance (B) of GSH after 1h, 5h, and 24h culture in [2,3,3-^2^H]-serine isotope enrichment. Panel C and D: isotopic enrichment (C) and ion abundance (D) of GSSG after 1h, 5h, and 24h of culture in [2,3,3-^2^H]-serine. Panel E and F: fractional isotopic enrichment (E) and ion abundance (F) of GSH after 48h culture in of [2,3,3-^2^H]-serine. Panel G-H: fractional isotopic enrichment (G) and ion abundance (H) of cystathionine after 48h of culture in [2,3,3-^2^H]-serine. All data represent mean ± S.D., sALS-1, n=3; control, n=3, *, P<0.05 (2-tailed Student’s t-test).

The biosynthesis of cytosolic GSH is tightly regulated by cysteine availability and the activity of γ-glutamylcysteinyl ligase (GCL)^13-15^. As shown in Fig 1F, cystathionine is synthesized by cystathionine β-synthase (CBS) via condensation of homocysteine with serine. Hence, the extent of [2,3,3-^2^H]-serine incorporation into cystathionine can serve to assess activity of the trans-sulfuration pathway. Surprisingly, we did not detect any serine incorporation into cystathionine at 1h, 5h and 24h after [2,3,3-^2^H]-serine supplementation, indicating that trans-sulfuration from homocysteine to *de novo* synthesized cysteine was insignificant during this period. One explanation for this apparent lack of *de novo* cysteine production is that cystine in the culture medium was ample to support GSH production over the 24h study period.

To assess whether activation of trans-sulfuration is indeed delayed by cysteine/cystine entry from the growth medium, we cultured cells with [2,3,3-^2^H]-serine for 48h, to allow for depletion of cystine in the medium. At this time, increased incorporation of serine-derived cysteine for GSH synthesis was observed in the sALS-1 group vs. control (as demonstrated by M+2 isotopologues, in addition to a predominant serine-derived M+1 glycine incorporation), while the total GSH pool was not significantly different (Fig 2E, F). In accord with utilization of serine for *de novo* cysteine synthesis, after 48 h we also observed increased incorporation of deuterium from serine into cystathionine in the sALS-1 subgroup vs. control (Fig 2G-H). Therefore, increased cysteine demand in the sALS subgroup, at least in part for GSH synthesis, resulted in accelerated trans-sulfuration of homocysteine to cysteine, concomitant with increased NADH dependent reduction of alpha-ketobutyrate to 2-hydroxybutyrate (Fig 1F). Notably, the cystathionine γ-lyase/hydrogen sulfide system was reported to be essential for maintaining cellular GSH^16^. Thus, upregulation of the trans-sulfuration pathway in sALS-1 can serve to maintain GSH levels for protection against oxidative stress.

### [U-^13^C]-glucose incorporation into GSH synthesis is increased in sALS-1

We next explored the relative utilization of glutamate for GSH synthesis. Glucose enters the TCA cycle as acetyl-CoA and contributes carbon atoms to produce α-ketoglutarate and its transamination product, glutamate. We compared the indirect contribution of [U-^13^C]-glucose-derived glutamate to GSH synthesis in sALS-1 and control lines. To this end, 5 mM [U-^13^C]-glucose was added to glucose-free DMEM medium and ^13^C incorporation into glutamate was quantified at 1h, 5h, 24h and 48h. At 1h, ^13^C enrichment of intracellular glucose approximated 100%, and incorporation into glutamate was observed at all time points predominantly as the M+2 isotopologue (Fig 3A-B). sALS-1 fibroblasts showed significantly increased GSH incorporation from glucose-derived glutamate (M+2) at 24h and 48h (Fig 3C, E). Notably, [U-^13^C]-glucose can also incorporate into serine, and serine-derived glycine can be further incorporated into GSH. However, under the cell culture conditions used, incorporation of glucose into glycine (M+1) was much less than into glutamate M+2 (Fig 3C-E). Despite increases in GSH incorporation from glutamate at 24h and 48h, the total GSH pool was similar in sALS-1 and control fibroblasts (Fig 3D, F), indicating that both GSH synthesis and consumption are increased in sALS-1 patient-derived fibroblasts. This interpretation is further strengthened by the accelerated incorporation of ^13^C into GSSG and the increased total GSSG pool in sALS-1 fibroblasts (Fig 3G-H). Together, these findings indicate an accelerated rate of GSH oxidation and enhanced NADPH consumption to provide antioxidant defense in fibroblasts from patients with the sALS-1 metabotype.

**Figure 3.**
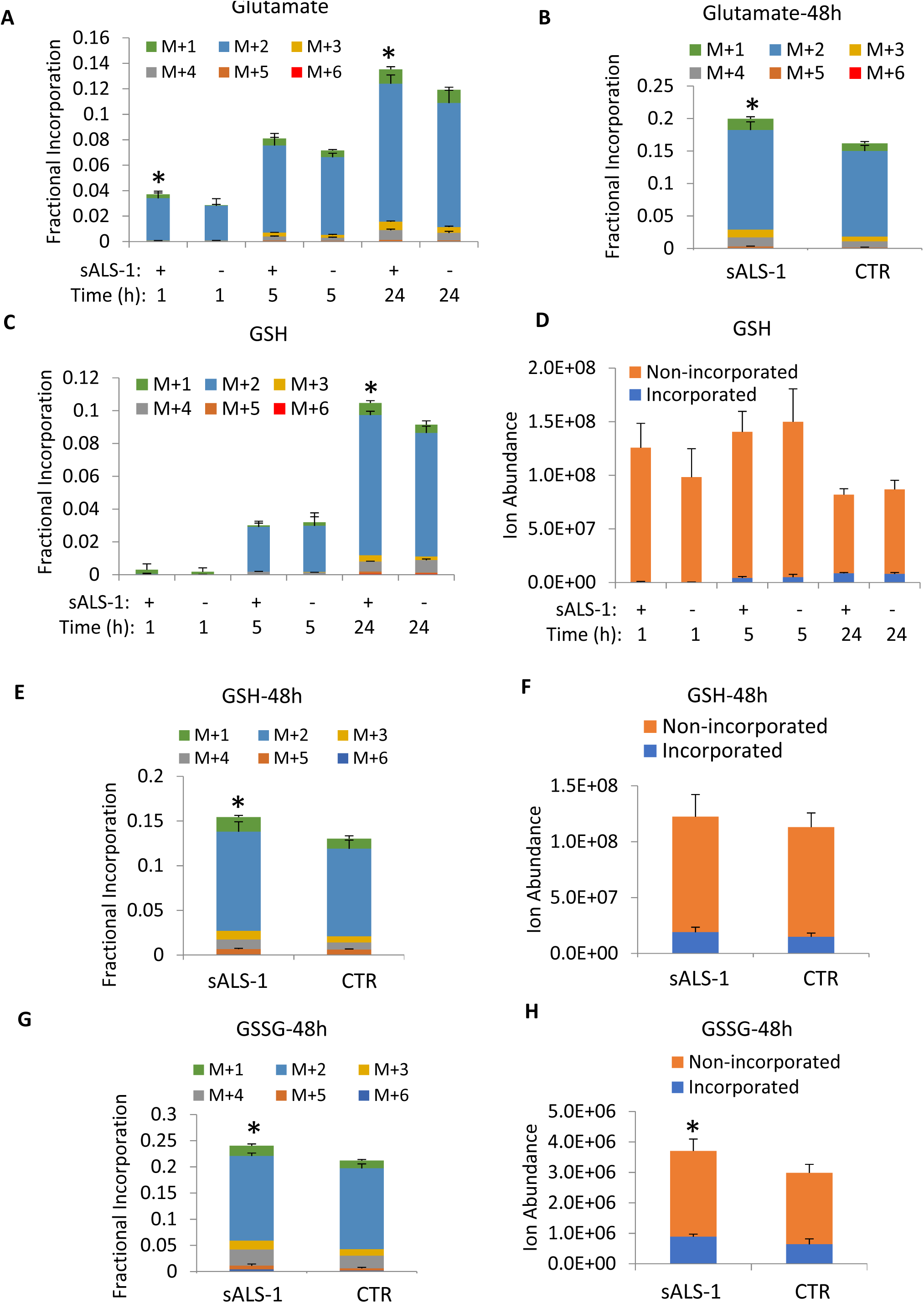
[U-^13^C]-glucose incorporation into GSH synthesis is increased in sALS-1. Panel A, B, C, E, G: Glutamate, GSH and GSSG fractional incorporation after 1h, 5h, 24h and 48h of [U-^13^C] glucose isotope enrichment. Panel D, F: ion abundance of GSH and GSSG after 1h, 5h, 24h and 48h of [U-^13^C] glucose isotope enrichment. All data represent mean ± S.D., sALS-1, n=3; control, n=3, *, P<0.05 (2-tailed Student’s t-test).

### [U-^13^C]-glucose incorporation into TCA cycle intermediates and nucleotides is accelerated in sALS-1

Stable isotope tracing studies also revealed that increased GSH synthesis in sALS1 was associated with an accelerated overall rate of glycolysis, TCA cycle, and additional linked metabolic pathways. We analyzed targeted and untargeted ^13^C tracing results from cells in [U-^13^C]-glucose medium at 1h, 5h and 24h. sALS-1 fibroblasts exhibited increased fractional incorporation of [U-^13^C]-glucose into lactate, TCA cycle intermediates, amino acids (aspartate and alanine), and C2 and C4 acyl-carnitines (Fig 4A-K). Interestingly, regardless of changes in total metabolite abundance (Fig S2-3), the rate of glucose incorporation of into each of the aforementioned species was increased at 24h. Additionally, we observed increased TCA cycle and cataplerotic products in an independent experiment, where cells were cultured for 48h in [U-^13^C]-glucose-supplemented medium (Fig S4). Notably, most TCA cycle intermediates exhibited <15% enrichment from ^13^C in glucose, except for citrate, which exhibited a 40% isotopic enrichment after 24h. This is consistent with the fact that rapidly proliferating fibroblasts operate a truncated TCA cycle, where citrate is diverted to the cytosol for oxidation to oxoglutarate by isocitrate dehydrogenase 1, concomitant with NADPH production and generation of acetyl-CoA for fatty acid synthesis^17^. Our data indicate that sALS-1 fibroblasts exhibit an accelerated TCA cycle flux, both oxidative and reductive, supportive of increased energy metabolism. This hypermetabolic phenotype is consistent with an increase in mitochondrial bioenergetics that was previously demonstrated generically for sALS vs. control fibroblasts^5, 9^

**Figure 4.**
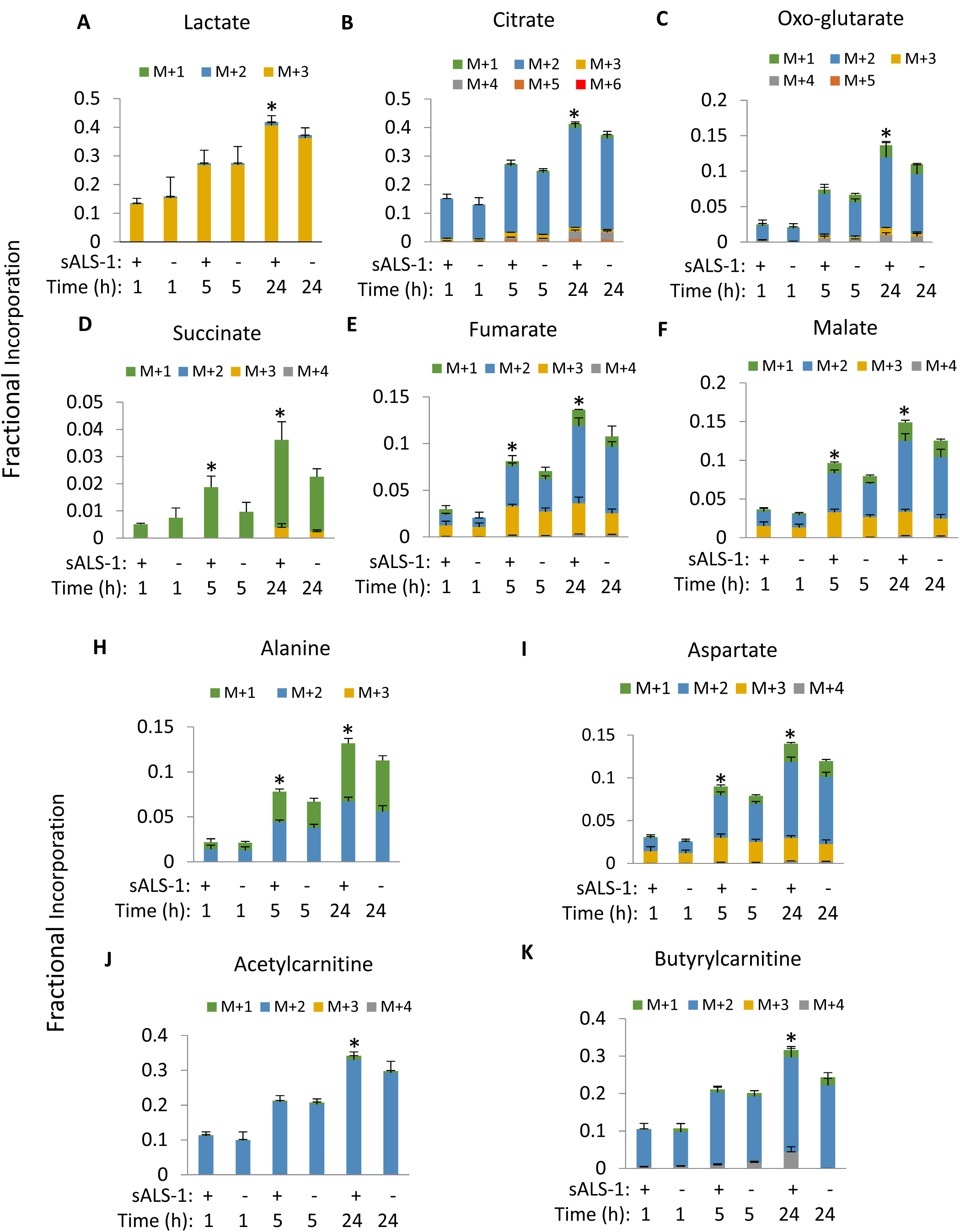
[U-^13^C]-glucose incorporation into lactate, TCA cycle intermediates and related metabolites is increased in sALS-1. Panel A-K: fractional incorporation after 1h, 5h, and 24h of [U-^13^C] glucose isotope enrichment. All data represent mean ± S.D., sALS-1, n=3; control, n=3, *, P<0.05 (2-tailed Student’s t-test). Note: Due to the overlapping of succinate M+2 with the reference ion (m/z 119.03) used for mass accuracy correction, the calculated M+2 incorporation in succinate was much smaller than the actual incorporated value. However, the relative incorporation of [U-^13^C] glucose to succinate is significantly increased in sALS-1 compared to the control under the same condition.

In accord with an accelerated TCA cycle, sALS-1 fibroblasts exhibited a significant increase in glucose incorporation into nucleotide triphosphate pools after 24h (mostly via the ribose M+5 isotopologue), total levels of nucleotide triphosphate remained unchanged (Fig 5 A-D, Fig S5 A-D). This can be explained by increased rates of both *de novo* nucleotide synthesis and nucleotide phosphate consumption. Notably, in sALS-1, 6-phosphogluconate, a precursor of NADPH, ribulose-5-phosphate, and ribose 5-phosphate production via the pentose phosphate pathway, showed a rapid fractional enrichment that reached 85% by 1h (Fig 5E). Increased fractional enrichment and decreased total intracellular levels (Fig S5E) similarly indicated a faster rate of 6-phosphogluconate consumption in sALS-1.

**Figure 5.**
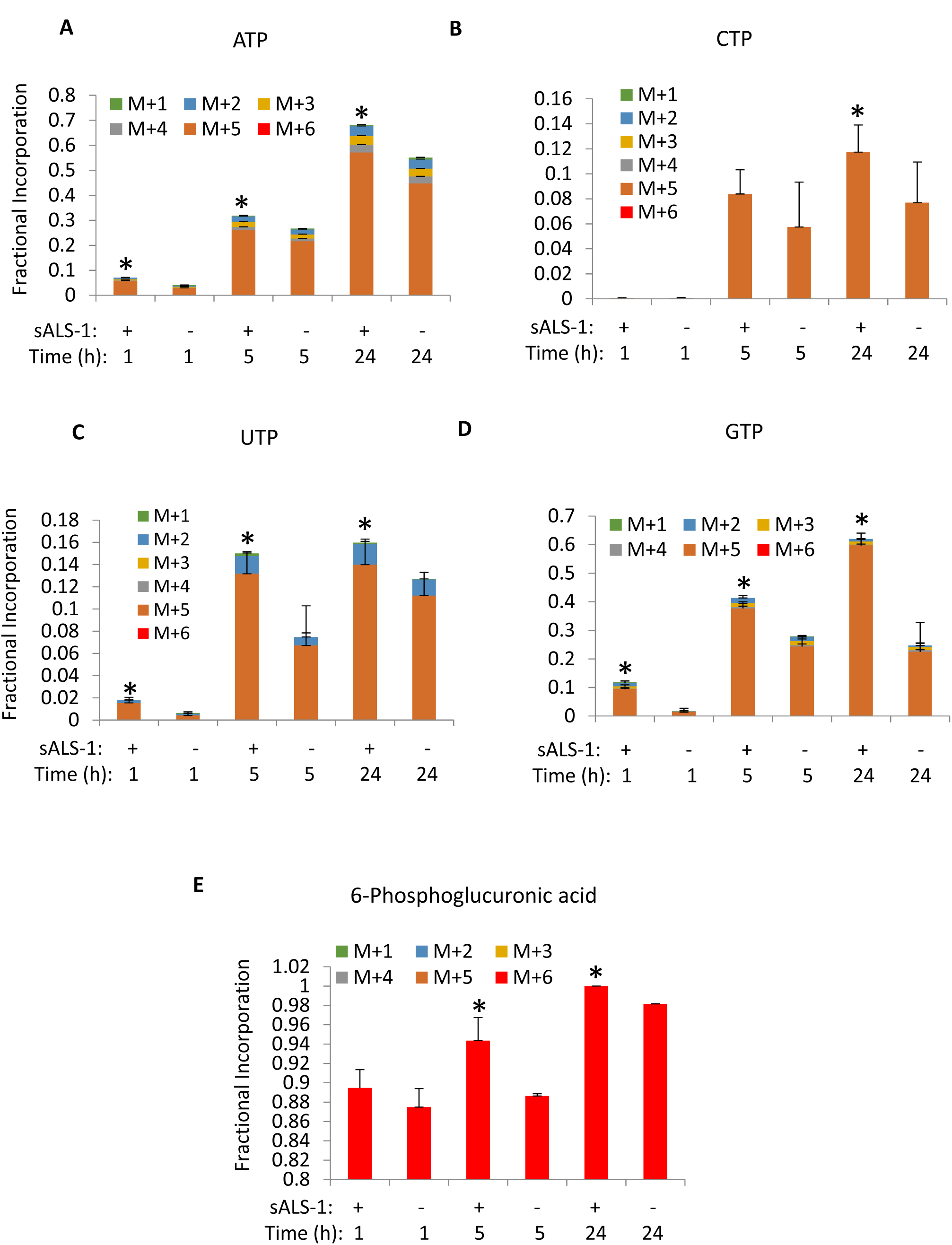
[U-^13^C] glucose incorporation into nucleotide triphosphates and pentose phosphate intermediate is increased in sALS-1. Panel A-K: fractional incorporation after 1h, 5h and 24h of isotope enrichment. All data represent mean ± S.D., sALS-1, n=3; control, n=3, *, P<0.05 (2-tailed Student’s t-test).

Taken together, these results indicate that sALS fibroblasts exhibit increased glucose influx into the TCA cycle, cataplerotic amino acid synthesis, accelerated GSH synthesis, and pentose phosphate pathway activity, providing antioxidant NADPH and ribose for increased nucleotide synthesis.

### Mitochondrial membrane potential is increased in sALS-1

We previously demonstrated that mitochondrial bioenergetics is perturbed in skin fibroblasts from a group of 171 sALS patients, compared with 98 control fibroblast cell lines^5^. Particularly, sALS fibroblasts showed a significant increase in mitochondrial membrane potential (MMP) as well as the ratio between MMP and mitochondrial mass (MMP:MM ratio). Since the sALS patient- and control-fibroblasts utilized in the present study were both included in the previously studied cohort, we performed meta-analysis of the prior bioenergetics data for sALS-1, relative to non-stratified sALS (denoted sALS-2) and control fibroblasts. Based on this analysis, sALS-1 with enhanced trans-sulfuration, but not sALS-2, exhibited lower MM and increased MMP relative to controls, and increased MMP:MM ratio relative to both controls and sALS-2 (Fig S6). Interestingly, metabolite profiling and glucose tracing of ribose incorporation into ATP synthesis revealed that the total intracellular ATP pools did not differ among these groups, despite a significantly increased ribose incorporation into ATP (Fig 5A, S5A). This finding is in agreement with the published reports using a chemiluminescent approach to quantify ATP levels^18^. Together, these data support the hypothesis that sALS-1 patients have increased bioenergetic demands^5^.

### Integrated transcriptomic and metabolomic pathway analysis reveals super-trans-sulfuration activation in sALS-1

We performed RNA microarray and microRNA analysis to investigate whether glucose and trans-sulfuration hypermetabolism can be explained by selective transcriptional changes in sALS-1 fibroblasts. Toward this end, transcripts and microRNA targets from 27 sALS fibroblasts were compared to those from 27 randomly selected control subjects. Of the 27 sALS patient cell lines studied, 11 belonged to the sALS-1 metabotype and 16 were sALS-2. Due to the comprehensive nature of gene annotation databases (mostly cancer and disease related), we first curated the searchable database to include candidate gene sets from previously published sALS studies, then identified targeted genes based on differentially-expressed transcripts and microRNA targets in sALS subgroups.

Using the curated database, we integrated transcriptome, microRNAome and metabolome data into metabolic pathways queried by KEGG, Wiki-pathways and BioCyc. The results identified the *super-trans-sulfuration pathway* to be among the top significantly enriched pathways in sALS-1 fibroblasts (p=2.3e-4 by Fischer’s exact test, Fig 6A). Note that sALS-1 fibroblasts showed a much smaller enrichment p-value than sALS-2, in accord with the trans-sulfuration pathway being significantly accelerated in sALS-1 relative to sALS-2 and control fibroblasts.

**Figure 6.**
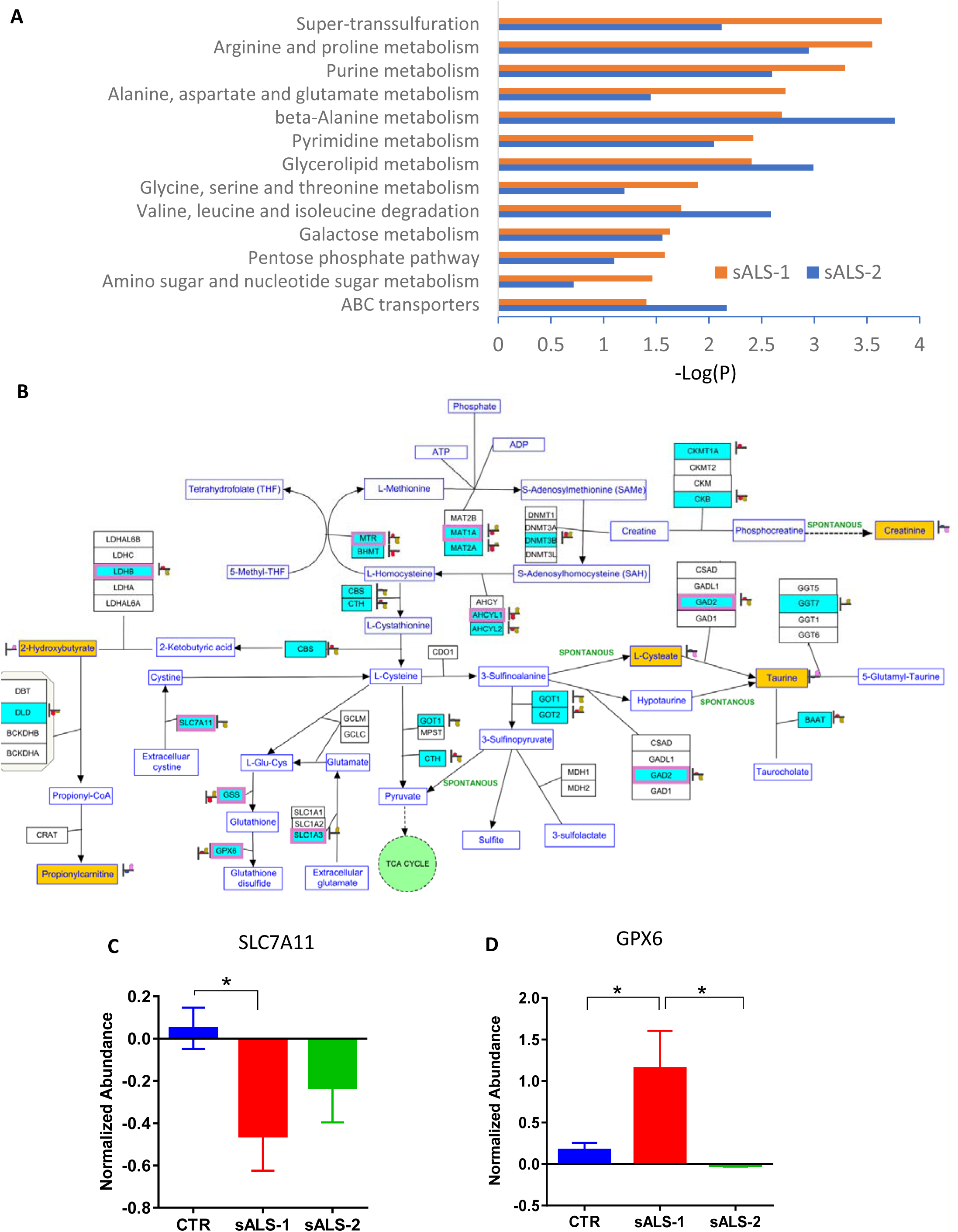
Pathway Integration of transcript and metabolite changes in sALS-1 and sALS-2 compared to controls. Panel A: Significant enrichments of integrated multiomic data into metabolic pathways in sALS-1 and sALS-2 compared to controls (P=2.3e-4, Fisher’s exact test). Differentially expressed messenger RNA (mRNA) and mircroRNA (miRNA) targets were obtained from 11 sALS-1, 16 sALS-2 and 27 age and gender matched controls. Differentially expressed intracellular metabolites were obtained from 18 sALS-1 relative to 18 controls. The metabolic pathways used for mapping were queried by KEGG, Wiki-pathways and Biocyc and imported into Agilent GeneSpring 14.9.1 for pathway analysis. Panel B: Visualization of significant mapping of the differentially expressed mRNA, miRNA targets and metabolites involved in the super-trans-sulfuration pathway in sALS-1 compared to control group. Green entities with purple boxes: differentially expressed mRNA (P<0.05, 2-tailed Student’s t test); green entities without purple box: differentially expressed miRNA targets (P<0.05, 2-tailed Student’s t test); yellow entities: intracellular metabolites. Heat strips next to entities represent normalized abundance in control (left heat strip) and sALS-1 (right heat strip). Panel C-D: Distinctive expression of mRNAs related to cysteine transport and glutathione oxidation in sALS-1 compared to sALS-2 and control group. (C) cystine-glutamate antiporter (SLC7A11) and (D) glutathione peroxidase 6 (GPX6); P<0.05, 2-tailed Student’s t test.

We next mapped the multiomics data (i.e. differentially expressed mRNA, microRNA targets and metabolites in sALS-1 compared to control) representing the trans-sulfuration pathway to concomitantly visualize differentially-expressed transcripts along with metabolites in sALS-1. Differentially-expressed mRNA transcripts (entities depicted in Fig 6B in green with purple boxes; P<0.05) and differentially expressed microRNA targets (entities depicted in Fig 6B in green without purple boxes; P<0.05) were not only detected in the methionine cycle and homocysteine to cysteine trans-sulfuration pathway, but also in the GSH and cysteine metabolic pathways (both synthesis and oxidation/degradation).

The cystine/glutamate antiporter (system xc-/SLC7A11) contributes importantly to GSH synthesis by transporting cystine into the cell. With cystine entry, SLCA11 concomitantly antiports glutamate, potentially leading to excitotoxicity. This dual action of SLCA11 may be highly relevant to ALS because of its key roles in both oxidative stress and excitotoxicity. SLC7A11 was previously reported to be dysregulated in ALS patients and animal models of disease^19-21^. Notably, significant downregulation of SLC7A11 was found in sALS-1, but not sALS-2, compared to controls (Fig 6C). Conversely, GPX6, an isoform of glutathione peroxidase involved in the oxidation of GSH to GSSG, was found to increase in sALS-1, but not sALS-2, relative to controls (Fig 6D). Decreased SLC7A11 and increased GPX6 expression could explain the enhanced trans-sulfuration in in sALS-1.

Taken together, sALS-1 case-derived fibroblasts show distinct transcriptional changes involved in cystine and glutamate transport, as well as GSH synthesis/oxidation, compared to sALS-2 and control fibroblasts.

### N-acetylcysteine selectively rescues metabolically-stressed sALS-1 fibroblasts

*To* further investigate whether cysteine availability and trans-sulfuration play are biologically relevant in sALS-1 fibroblasts, we compared the effect of N-acetylcysteine (NAC) supplementation on cell proliferation under metabolic stress conditions, where galactose was substituted for glucose as a bioenergetic fuel in the fibroblast culture media. Whereas the production of pyruvate via glycolytic metabolism of glucose yields 2 net molecules of ATP, the production of pyruvate via glycolytic metabolism of galactose yields no net ATP. Thus, galactose medium forces cells to rely on mitochondrial oxidative phosphorylation (OXPHOS) and has been used to test fibroblasts for oxidative defects^22^.

Compared to sALS-2 and control fibroblasts, sALS-1 metabotype fibroblasts exhibit upregulated oxidative phosphorylation in glucose-containing medium (Fig S6 and Figs 4-5), and predictably cannot upregulate OXPHOS to a further extent in galactose-containing medium. As shown in Fig 7A, sALS-1 fibroblasts showed significantly impaired viability when grown in galactose medium at 24 h, and this growth defect was completely rescued by NAC supplementations. In contrast, sALS-1 cell viability in glucose-containing medium was not different from control and sALS-2 fibroblasts, and NAC supplementation had no effect on cell viability in any of the groups in glucose medium (data not shown).

**Figure 7.**
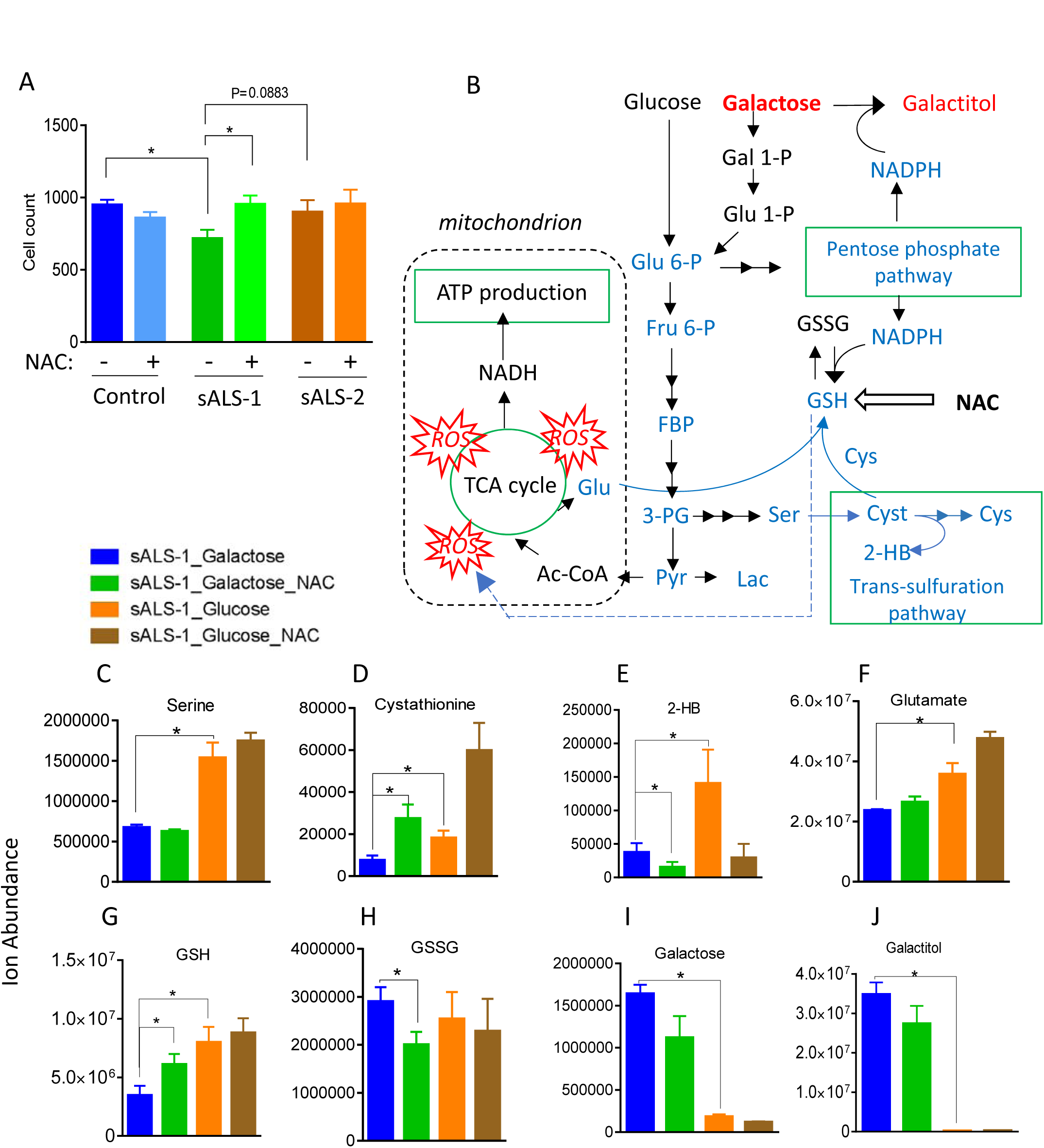
NAC selectively rescues sALS-1 fibroblasts exposed to metabolic stress. (A) sALS-1 fibroblasts show a distinct loss of viability phenotype in galactose medium, rescued by 24h 2mMNAC supplementation, compared to controls and sALS-2 group. n=18-40 from 3 cell lines per group, *, P<0.05, 2-tailed Student’s t-test. (B) Schematic depicting key metabolites in galactose glycolysis, TCA cycle, pentose phosphate pathway and trans-sulfuration pathway that contribute to galactose induced oxidative stress and subsequent NAC rescue in sALS-1. Glu 6-P, glucose 6-phosphate; Gal 1-P, galactose 1-phosphate; Glu 1-P, glucose 1-phosphate; Fru 6-P, fructose 6-phosphate; FBP, Fructose 1,6-bisphosphate; 3PG, 3-phosphoglycerate; Pyr, pyruvate; F1P, fructose-1-phosphate; Lac, lactate; Ac-CoA, Acetyl-CoA; Glu; Glutamate; Ser, serine; Cyst, cystathionine; 2-HB, 2-hydroxybutyrate; Cys, cysteine. Blue denotes decreased and red denotes increased metabolites and associated pathways in galactose compared to glucose medium. (C-J) key metabolite changes sALS-1 comparing galactose and glucose media with and without NAC treatment. n=8-9 from 3 cell lines per group, *, P<0.05, 2-tailed Student’s t-test.

Besides forcing OXPHOS, galactose is known to induce oxidative stress^23-25^. Consistently, we found that key metabolites related to antioxidant defense are altered in galactose-relative to glucose-containing sALS-1 fibroblast growth media (Fig 7, B-J). Serine, cystathionine and 2-hydroxybutyrate, the precursor and products of the trans-sulfuration pathway, were significantly decreased in galactose medium compared to glucose medium (Fig 7C-E), indicating the capacity of sALS-1 cells to compensate for heightened oxidative stress in galactose-containing medium by upregulating trans-sulfuration was compromised. Diminished synthesis of serine from glycolysis and synthesis of glutamate from the TCA cycle (Fig 7F) may underlie the observed decrease in GSH synthesis (Fig 7G), while GSSG remains unchanged (Fig 7H). Additionally, a decreased rate of glucose-6-phoshate production in galactose-containing medium results in decreased pentose phosphate flux and NADPH production. A slowed rate of glycolysis also leads to galactose build-up, and subsequent activation of galactose reduction to galactitol, which consumes NADPH (Fig 7B, I-J). Notably, the combination of increased galactose reduction to galactitol and decreased NADPH production from pentose phosphate pathway impairs the recycling of GSH from GSSG. NAC supplement effectively prevented galactose-induced oxidative stress, decreased GSSG, increased GSH synthesis, and enhanced cell viability (Fig 7A, G-H). These results support the view that that sALS-1 fibroblasts are operating OXPHOS and trans-sulfuration at near capacity in glucose-containing medium and are incapable of further upregulation when challenged with an oxidative stressor. Our findings indicate that NAC supplementation provides sALS-1 fibroblasts with sufficient antioxidant capacity to enable metabolic diversion from trans-sulfuration to energy generation.

### sALS-1 patient plasma displays altered trans-sulfuration pathway metabolites

To identify novel disease biomarkers, we explored if sALS-1 patient plasma exhibits indications of altered metabolism, in accord with findings in fibroblasts form this patient subset. To test this possibility, plasma samples from the same 18 sALS-1 cases that yielded fibroblasts with aberrant trans-sulfuration/hyper-metabolism phenotype were compared to plasmas from 20 age/sex-matched controls. Targeted and untargeted metabolite profiling detected >1000 metabolite features, of which 85 were found to be differentially-abundant in sALS-1 cases, compared to controls (Fig 8A). PCA analysis revealed that plasma metabolite abundances from sALS-1 cases cluster and separate from controls based on levels of these 85 metabolites (Fig 8B). Notably, the metabolite differences included decreased levels of some amino acids (glutamine, glutamate, methionine, tyrosine) and creatinine, along with increased levels of taurine-related species (i.e., taurine and glutamyl-taurine, bile acids and 2-hydroxybutyrate), purine metabolites, phospholipids, ceramides and fatty acid amides. Two species that were initially unknown and found to be present in the majority of sALS-1 cases were subsequently identified as Riluzole and its primary metabolite, hydroxyriluzole glucuronide. Mapping all identified differentially abundant sALS-1 plasma metabolites to KEGG pathways, revealed taurine/hypotaurine metabolism as the pathway with highest impact score and most significant p-value (Fig 8C), consistent with the results from sALS-1 fibroblast studies. Notably, increased plasma taurine and 2-hydroxybutyrate (both trans-sulfuration pathway intermediates), along with decreased creatinine (methionine cycle-related methylation product; Fig 1F), exhibited similar alterations as in fibroblasts, suggesting a systemic metabolic perturbation in sALS-1 patients, and the potential for effective use of plasma profiling for facile clinical recognition of the sALS-1 metabotype.

**Figure 8.**
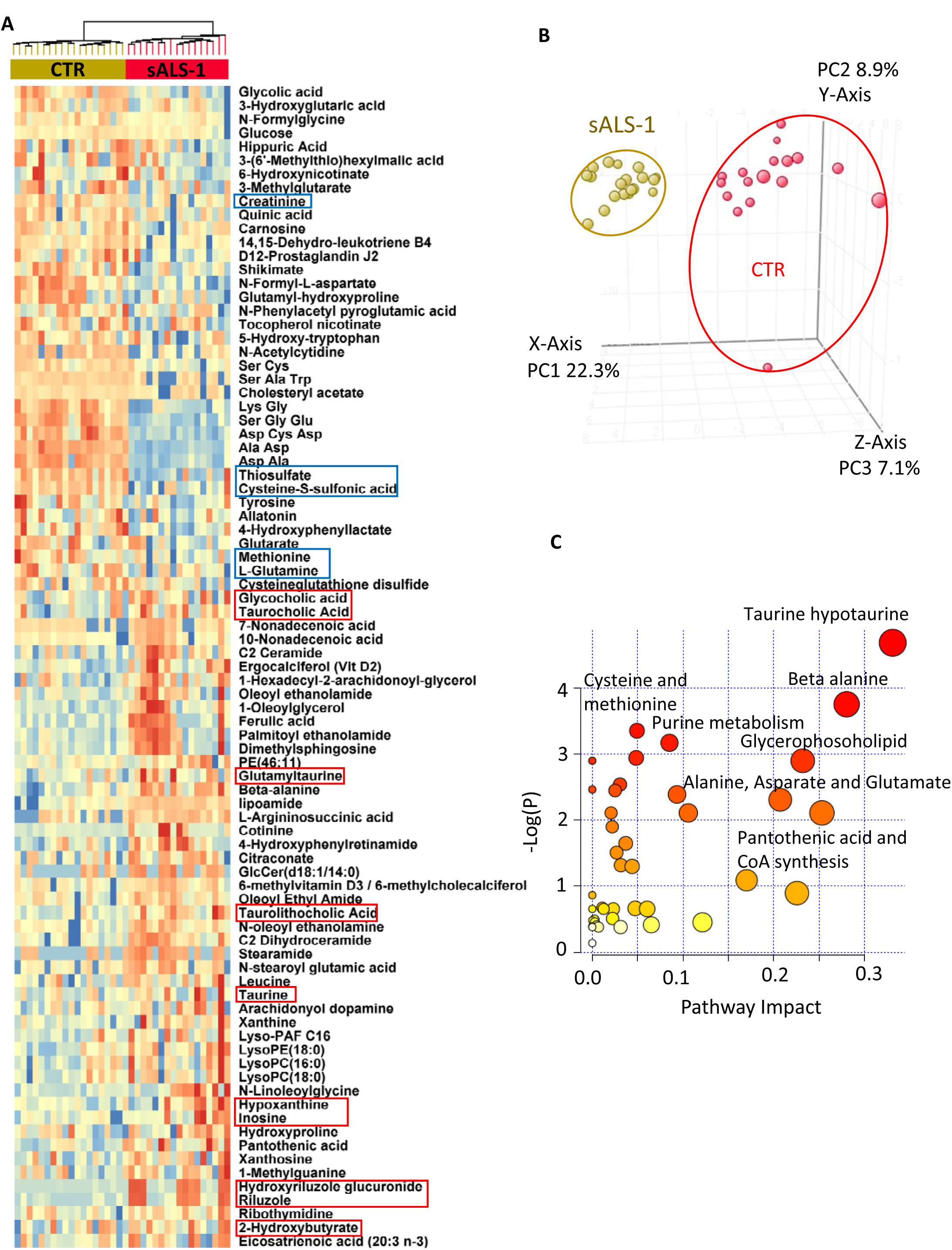
sALS-1 showed distinct plasma metabolite profiles from controls and indications of altered taurine and hypotaurine metabolic pathway. Panel A: heatmap of differential plasma metabolites in sALS-1 compared to controls. Metabolites related to altered taurine and trans-sulfuration pathway, amino acids, lipid metabolism and purine nucleosides were highlighted in red (upregulated) and blue (downregulated) boxes. ALS drug Riluzole and its drug metabolite hydroxyriluzole glucuronide were also highlighted. Panel B: PCA score plot showing the separation of sALS-1 from controls based on the 85 differentially expressed plasma metabolites (sALS-1, n=18; controls, n=20). Panel C: Mapping the differentially expressed plasma metabolites into KEGG pathway using Pathway analysis module of MetaboAnalyst 3.0 recognizes taurine and hypotaurine in the trans-sulfuration pathway as the most significant pathway impact and lowest matching P-value.

We then asked whether sALS-1 could be distinguished from sALS-2 patients by the clinical characteristics described in Table 1. Based on the available clinical data for 18 sALS-1 and 55 sALS-2, sALS-1 patients did not show significant difference in age at the onset of symptoms, ALSFRS-R (ALS functional rating scale–revised) at the time of biopsy, rate of functional decline, FVC (Forced vital capacity,%), and BMI(body mass index), relative to sALS-2. However, sALS-1 and sALS-2 exhibited different frequencies distribution for the site of initial disease onset (Fig S7). sALS-1 cases presented with a higher frequency of lumbosacral onset (50%), while sALS-2 had >50% cervical onset. These data raise an interesting question whether patient metabotype plays a role in the site of disease initiation, which in some studies has been shown to play a role in disease prognosis^26^.

## Discussion

In an effort to identify metabolic biomarkers for stratification of sALS patients, we applied untargeted metabolite profiling to dermal fibroblasts. This effort confidently identified a subgroup of sALS fibroblasts (i.e., sALS-1) with a distinct trans-sulfuration pathway upregulation. Importantly, we identify this metabotype also in cognate plasma from sALS-1 cases and show that this subgroup also exhibits perturbed mRNAs for trans-sulfuration/cysteine metabolism. We further demonstrate that the altered metabolic pathways in sALS-1 are linked to increased GSH synthesis and accelerated glucose metabolism. Notably, these metabolic alterations are associated with impaired cell viability under oxidative stress, which is highly responsive to supplementation with the antioxidant, NAC. This interplay between glucose hypermetabolism and increased antioxidant demand raises the possibility that upregulated antioxidant defenses serve to meet hypermetabolic demands in sALS-1 patients and define the disease phenotype.

It is known that a subset of ALS patients exhibits approximately 10% more brain resting energy expenditure than healthy individuals^27-29^. Overall, hypermetabolism is a feature of more than half of the ALS population and was recently shown to be a deleterious prognostic factor. Patients with hypermetabolism have a 20% worse prognosis than those that do not exhibit the hypermetabolism phenotype^30, 31^. Until now, hypermetabolism in ALS has mostly been considered in relation to glucose metabolism. Indeed, ^18^F-FDG-PET imaging is widely applied clinically to assess and discriminate CNS differences in glucose uptake that distinguish hypermetabolism in both familial and sALS patients^27, 32, 33^. Enhanced ^18^F-FDG-PET uptake correlates with cognitive impairment in ALS patients and provides a convenient means to identify altered metabolic activity^32^.

Using stable isotope tracing, we demonstrate a selective enhancement of trans-sulfuration pathway flux in sALS-1 fibroblasts, associated with accelerated rates of GSH synthesis and glucose metabolism. Moreover, increased intracellular and plasma levels of 2-hydroxybutyrate correlate with increased GSH synthesis and increased taurine abundance in the newly-identified sALS-1 subgroup. Since limited cystine uptake by fibroblasts appears to accelerate the conversion of homocysteine to cysteine via the trans-sulfuration pathway, fueling production of GSH, we surmise that sALS-1 patients have greater demand for anti-oxidant synthesis. We propose that increased glucose utilization and GSH synthesis are respectively a cause and consequence of oxidant stress in the sALS-1 fibroblast metabotype, found in approximately 25% (18/77) of sALS cases studied. In accord with this finding, a recent report correlated *in vivo* ALS staging with glucose metabolism patterns and demonstrated that four neuropathological stages of ALS correlate with discriminative regional brain glucose metabolism patterns that associate with both disease duration and forced vital capacity^34^. Therefore, sALS-1 cases are likely to comprise a heterogenous spectrum of glucose hypermetabolism phenotypes, with a distinct metabotype tuned by cysteine availability to enable antioxidant defenses for evasion of oxidative stress.

The finding that similar changes in trans-sulfuration metabolites occur in both fibroblasts and plasma from sALS-1 cases is intriguing, as it may provide an opportunity for facile stratification of sALS cases based on plasma biomarkers. Remarkably, changes in plasma levels 2-hydroxybutyrate, taurine, amino acids and creatinine have been previously reported in ALS patients^35-38^. As mentioned above, 2-hydroxybutyrate derives from 2-ketobutyrate, produced from cystathionine in the trans-sulfuration pathway via a NADH-dependent reduction catalyzed by lactate dehydrogenase B (Fig 1F). Elevated levels of plasma 2-hydroxybutyrate and 2-ketobutyrate have previously been shown in several studies involving different cohorts of ALS cases and ascribed to disease-associated oxidative stress^35-37^. Moreover, decreased plasma creatinine has previously been suggested to be an ALS risk factor^39, 40^. Besides taurine, bile acids (e.g., cholic acid) and bile acid conjugates (e.g., taurocholic acid) were found to increase in sALS-1 plasma (Fig 8A). Bile acids play a neuroprotective role in a diverse spectrum of age-related neurodegenerative disorders^41^. There are two primary bile acids produced by the liver in humans: cholic acid and chenodeoxycholic acid. These primary bile acids can undergo conjugation with glycine or taurine prior to secretion in the bile as glycocholic acid, taurocholic acid, glycochenodeoxycholic acid, and taurochenodeoxycholic acid. Therefore, upregulated plasma levels of taurine and taurocholic acid could represent an adaptive defense mechanism that opposes heightened oxidative stress.

In summary, we describe the first stratification of a subgroup of sALS skin fibroblasts based on metaboytpe and identify a distinct patient cohort characterized by increased trans-sulfuration and glucose hypermetabolism. We show that the observed metabolite changes are similarly detected in sALS-1 plasma and find that sALS-1 fibroblasts are uniquely sensitive to metabolic stress and selectively protected by supplementation with the antioxidant NAC. Therefore, we infer that facile plasma profiling could be employed to confidently stratify sALS cases, recognizing the sASL-1 subtype for targeted benefit using anti-oxidant therapies. We hypothesize that the metabolic rewiring that results in upregulation of the trans-sulfuration pathway in these patients arises as an adaptation to limiting cysteine and hence GSH, potentially triggered by hypermetabolism that accelerates GSH oxidation. Knowledge obtained from metabotypic stratification of skin-derived fibroblasts from sALS-1 cases may provide a much-needed first step toward developing precision medicines for therapy of sALS patients.

## Materials and Methods

### Reagents

LC-MS grade acetonitrile (ACN), isopropanol (IPA) and methanol (MeOH) were purchased from Fischer Scientific. High purity deionized water (ddH2O) was filtered from Millipore (18 OMG). OmniTrace glacial acetic acid and ammonium hydroxide were obtained from EMD Chemicals. [U-13C] glucose, [2,3,3-^2^H]-serine were purchased from Cambridge Isotope Laboratory. Ammonium acetate and all other chemicals and standards were obtained from Sigma Aldrich in the best available grade.

### Cell culture

A total of 77 sALS and 45 control fibroblast cell lines were obtained from cases and propagated as previously described (Konrad et al. Mol Neuorodeg 2017). 75,000 cells/well for each line were plated in duplicate 6-well plates and cultured in DMEM medium containing 5 mM glucose and 4 mM glutamine, 10% FBS,1% of 100X antibiotic/antimycotic (Ab+F; which contains sterile-filtered 10,000 units penicillin, 10 mg streptomycin and 25 μg amphotericin B per mL, and 2.5 µg/ml Plasmocin). Cells were harvested at 80% confluency and extracted for LC/MS metabolomic analysis.

### Metabolite extraction

Each cell line was cultured and initially extracted as two biological replicates, for independent LC/MS metabolomic data acquisition. Cells were washed twice with ice-cold PBS, followed by metabolite extraction using −70°C 80:20 methanol:water (LC-MS grade methanol, Fisher Scientific). The tissue–methanol mixture was subjected to bead-beating for 45 sec using a Tissuelyser cell disrupter (Qiagen). Extracts were centrifuged for 5 min at 5,000 rpm to pellet insoluble material and supernatants were transferred to clean tubes. The extraction procedure was repeated two additional times and all three supernatants were pooled, dried in a Vacufuge (Eppendorf) and stored at −80°C until analysis. The methanol-insoluble protein pellet was solubilized in 0.2 M NaOH at 95°C for 20 min and protein was quantified using a BioRad DC assay. On the day of metabolite analysis, dried cell extracts were reconstituted in 70% acetonitrile at a relative protein concentration of 1 µg/ml, and 4 µl of this reconstituted extract was injected for LC/MS-based untargeted metabolite profiling.

These 244 patient-derived skin fibroblast extracts (122 samples with 2 biological replicates) were analyzed by LC-QTOF metabolomics profiling in random sequence. To adjust for day-to-day and batch-to-batch LC/MS instrument drift, metabolite stability, and other experimental factors that may contribute to systematic error, metabolite measurements were normalized to one another using flanking quality control (QC) samples run at intervals of every 6 injections. These QC samples were prepared from a pool of all samples and this normalization procedure enabled the comparative analysis of data acquired over a 1-month collection period.

Plasma metabolites were extracted by addition to 1 part plasma to 20 parts 70% acetonitrile in ddH2O (vol:vol). The mixture was briefly vortexed and then centrifuged for 5 min at 16,000 × g to pellet precipitated proteins. An aliquot of the resulting extract (3 μl) was subjected to untargeted metabolite profiling using, applying both positive and negative ion monitoring MS.

### Untargeted metabolite profiling by LC/MS

Cell extracts were analyzed by LC/MS as described previously^42, 43^, using a platform comprised of an Agilent Model 1290 Infinity II liquid chromatography system coupled to an Agilent 6550 iFunnel time-of-flight MS analyzer. Chromatography of metabolites utilized aqueous normal phase (ANP) chromatography on a Diamond Hydride column (Microsolv). Mobile phases consisted of: (A) 50% isopropanol, containing 0.025% acetic acid, and (B) 90% acetonitrile containing 5 mM ammonium acetate. To eliminate the interference of metal ions on chromatographic peak integrity and electrospray ionization, EDTA was added to the mobile phase at a final concentration of 6 µM. The following gradient was applied: 0-1.0 min, 99% B; 1.0-15.0 min, to 20% B; 15.0 to 29.0, 0% B; 29.1 to 37min, 99% B. Raw data were analyzed using MassHunter Profinder 8.0 and MassProfiler Professional (MPP) 14.9.1 software (Agilent technologies). Mann Whitney t-tests (p<0.05) were performed to identify significant differences between groups.

### Metabolite Structure Specification

To ascertain the identities of differentially expressed metabolites (P<0.05), LC/MS data was searched against an in-house annotated personal metabolite database created using MassHunter PCDL manager 7.0 (Agilent Technologies), based on monoisotopic neutral masses (<5 ppm mass accuracy) and chromatographic retention times. A molecular formula generator (MFG) algorithm in MPP was used to generate and score empirical molecular formulae, based on a weighted consideration of monoisotopic mass accuracy, isotope abundance ratios, and spacing between isotope peaks. A tentative compound ID was assigned when PCDL database and MFG scores concurred for a given candidate molecule. Tentatively assigned molecules were verified based on a match of LC retention times and/or MS/MS fragmentation spectra for pure molecule standards contained in a growing in-house metabolite database.

### Stable isotope tracing of [U-^13^C] glucose and [2,3,3-^2^H] serine

An in-house untargeted stable isotope tracing (USIT) workflow was employed using Agilent metabolite profiling software MassHunter Qualitative Analysis 7.0, MassProfinder 8.0 and MPP 14.9.1. Labelled metabolites were identified on the basis of differential abundance in cells cultured with supplemental heavy isotope-labelled metabolite vs. natural (light) isotope, as described previously^44, 45^. Selected metabolites were identified on the basis of previously curated isotopologues. Notably, the USIT workflow calculates and corrects for the natural abundance of ^13^C and ^2^H isotope in samples.

### Microarray analysis

Total RNA (combined mRNA and miRNA) from skin fibroblasts was extracted from 27 sALS patients and 27 controls using Trizol agent (Invitrogen). Total RNA was further extracted using the Agilent Absolutely RNA miRNA Kit. The total RNA concentration and RNA sample integrity were verified by a Bio-Analyzer 2100 (Agilent Technologies, Waldbronn, Germany). The quality of isolated RNA was determined using an Agilent 2200 TapeStation system and Bioanalyzer with mRNA labeling and microarray processing were performed according to the manufacturers recommendations. miRNA labeling was done using an Agilent miRNA Complete Labeling and Hyb Kit with gene expression and miRNA data extracted using Agilent Feature Extraction Software.

Extracted mRNA and miRNA data were analyzed using the respective workflows in GeneSpring GX 14.9.1 (Agilent Technologies, CA). Signal intensities for each mRNA probe was normalized to 75th percentile values with baseline transformation for gene expression analysis. Separately, miRNA data was normalized to 90th percentile and baseline transformed for miRNA analysis. The differential miRNA list (P<0.05, FC>1.2) was used to identify the gene targets based on a target prediction database incorporated in GeneSpring GX (e.g. TargetScan, PicTar, microRNA.org). The differentially expression mRNA was combined with validated miRNA targets and metabolites for integrated multi-Omics pathway analysis using databases linked to KEGG, Wiki pathways and Biocyc.

### Statistics

All values are averages of at least three independent measurements. Error bars indicate standard deviation (S.D.) or standard error of the mean (S.E.M.). Statistically significant differences between two groups were estimated by unpaired two-tailed Student’s test with significance set at p<0.05.

## Acknowledgements

Funding: NIH R01NS093872 (GM, SG, and LS), R01NS062055 (GM). We thank Dr. Hiroshi Mitsumoto (Columbia University) and the COSMOS initiative for providing the fibroblast and plasma samples utilized in this work.

## Author contributions

DS, BIS, RRC performed metabolite extraction and targeted metabolite analysis, DR and SMF analyzed the transcriptomics data. CK, KB, and HK performed cell culture and the fibroblast bioenergetics analysis. ELC and LC contributed to the data interpretation and concepts presented; QC, GM and SSG conceived the experiments, analyzed the data and wrote the initial manuscript draft for editing by co-authors.

## Supplemental Figure legends

**Figure S1.**
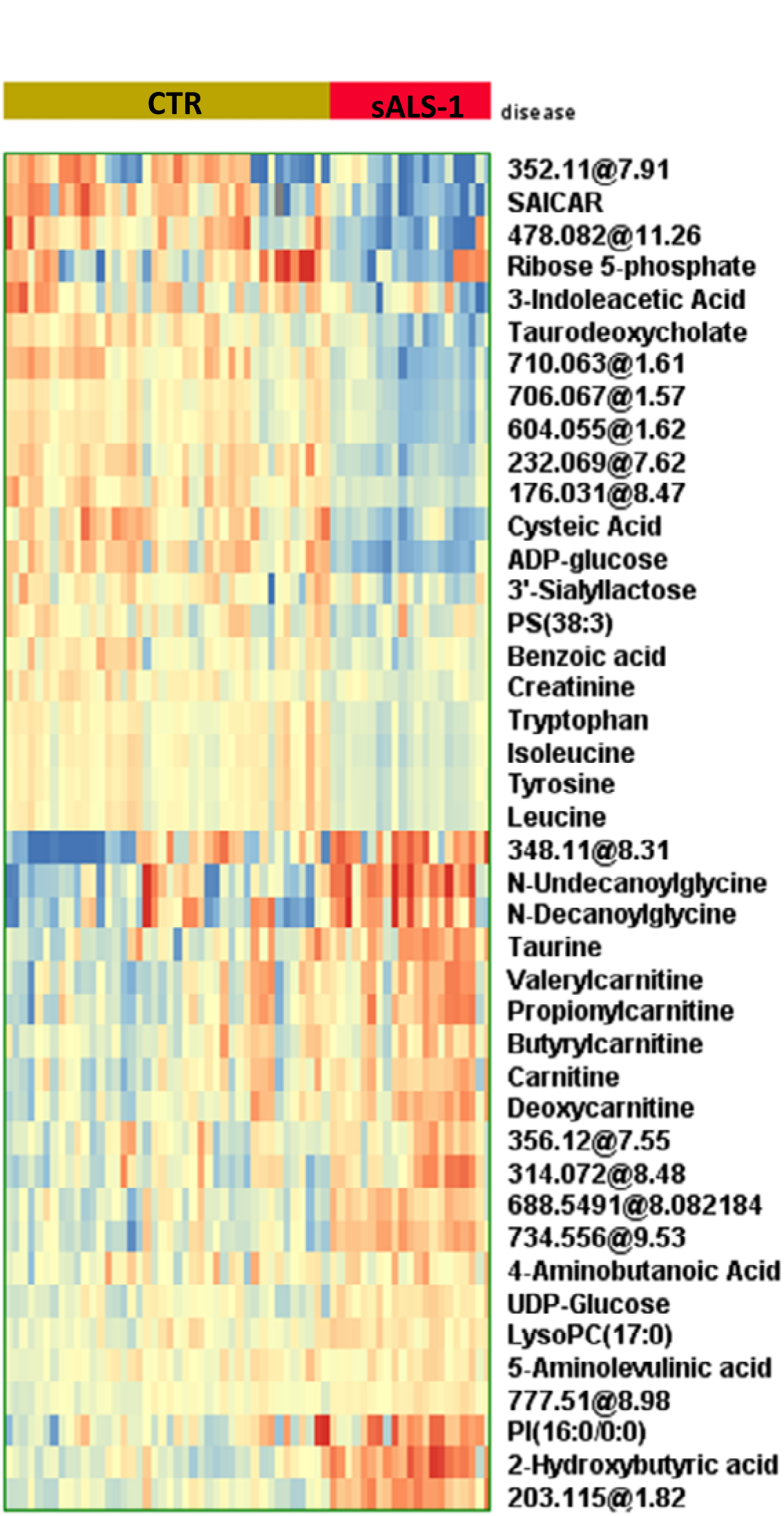
Heatmap depiction of 42 differential metabolites in 18 sALS-1 subgroup compared to 45 control. Unknown metabolites were expressed as accurate mass and retention time. Blue denotes downregulation and red denotes upregulation.

**Figure S2.**
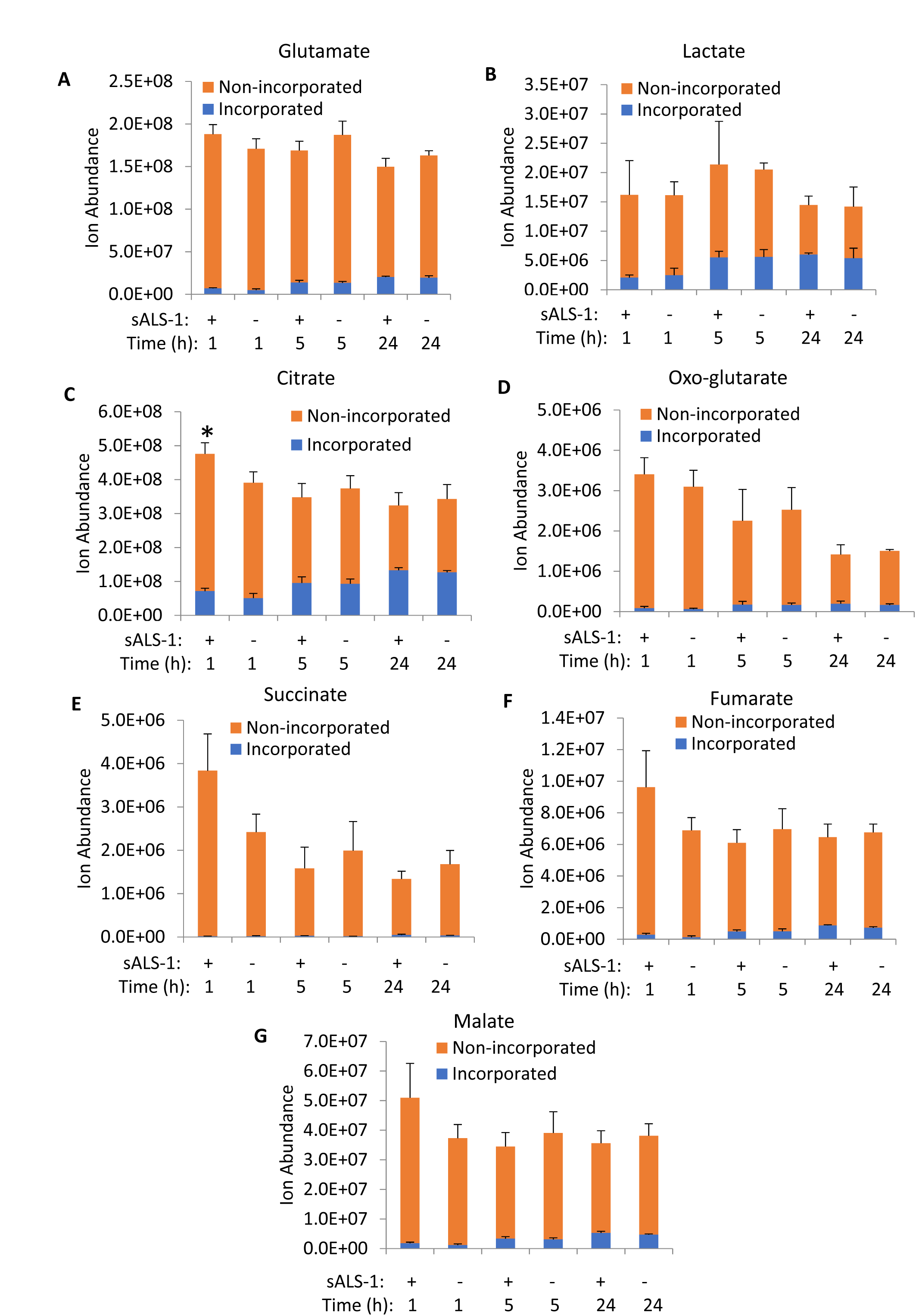
Relative abundance of glutamate, lactate and TCA cycle intermediates after 1h, 5h, and 24h of [U-^13^C] glucose enrichment. Data represent mean ± S.D., sALS-1, n=3; control, n=3, *, P<0.05 (2-tailed Student’s t-test).

**Figure S3.**
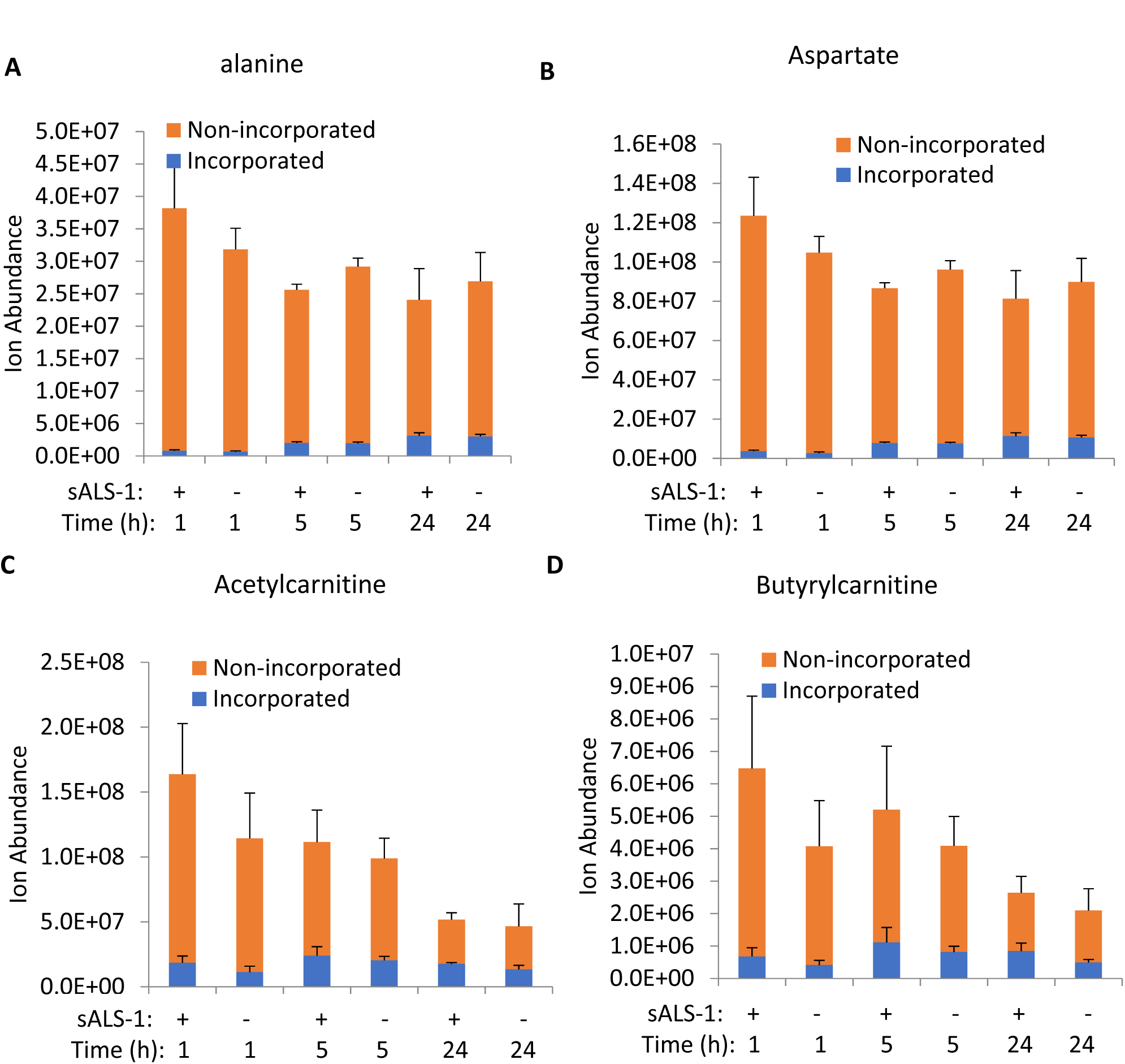
Relative abundance of amino acids and acylcarnitines after 1h, 5h, and 24h of [U-^13^C] glucose enrichment. Data represent mean ± S.D., sALS-1, n=3; control, n=3.

**Figure S4.**
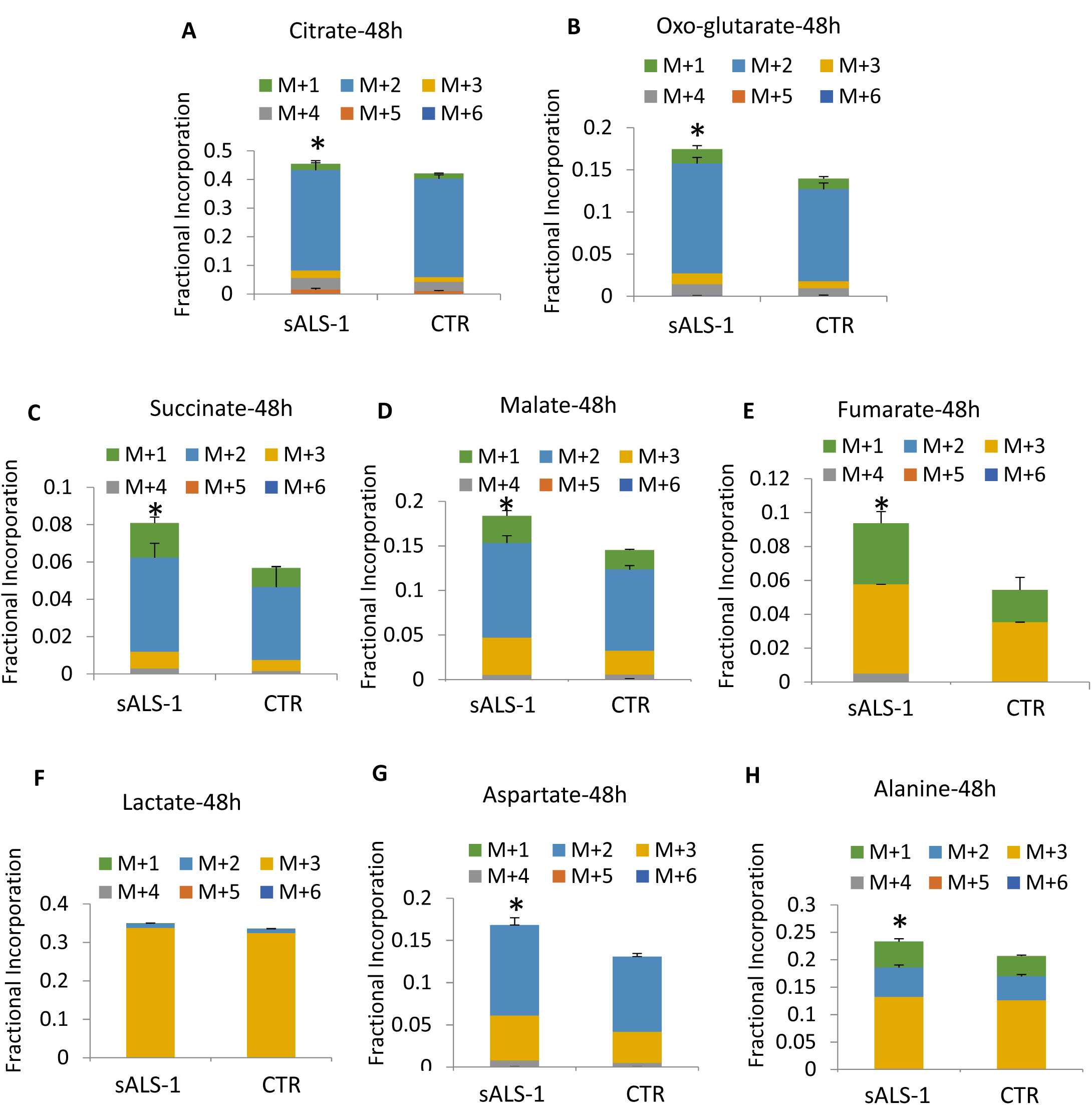
[U-^13^C]-glucose incorporation into lactate, TCA cycle and related intermediates is increased in sALS-1 after 48h isotope enrichment. Data represent mean ± S.D., sALS-1, n=3; control, n=3, *, P<0.05 (2-tailed Student’s t-test).

**Figure S5.**
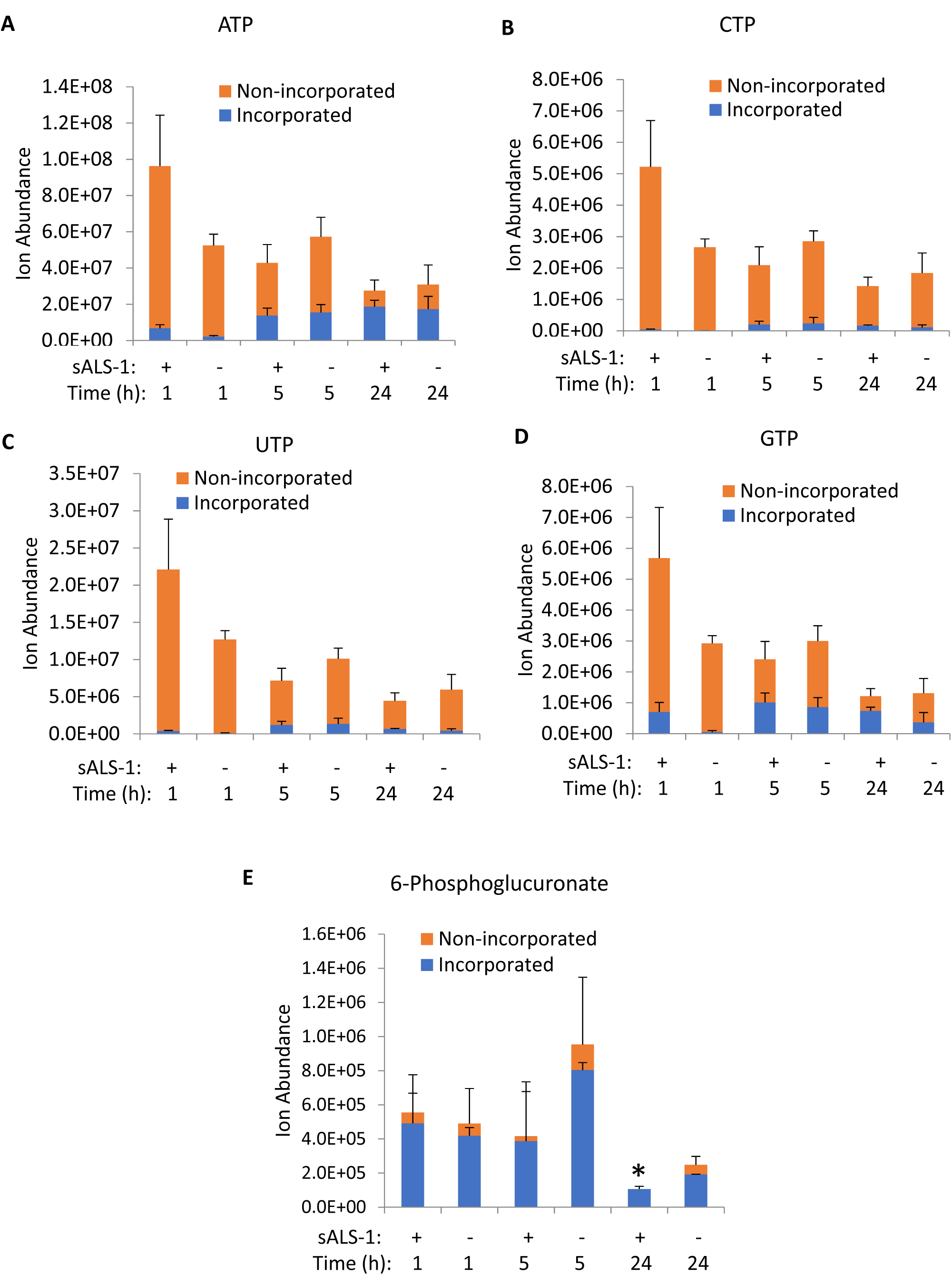
Relative abundance of nucleotide triphosphates and pentose phosphate intermediate after 1h, 5h, and 24h of [U-^13^C] glucose enrichment. Data represent mean ± S.D., sALS-1, n=3; control, n=3, *, P<0.05 (2-tailed Student’s t-test).

**Figure S6.**
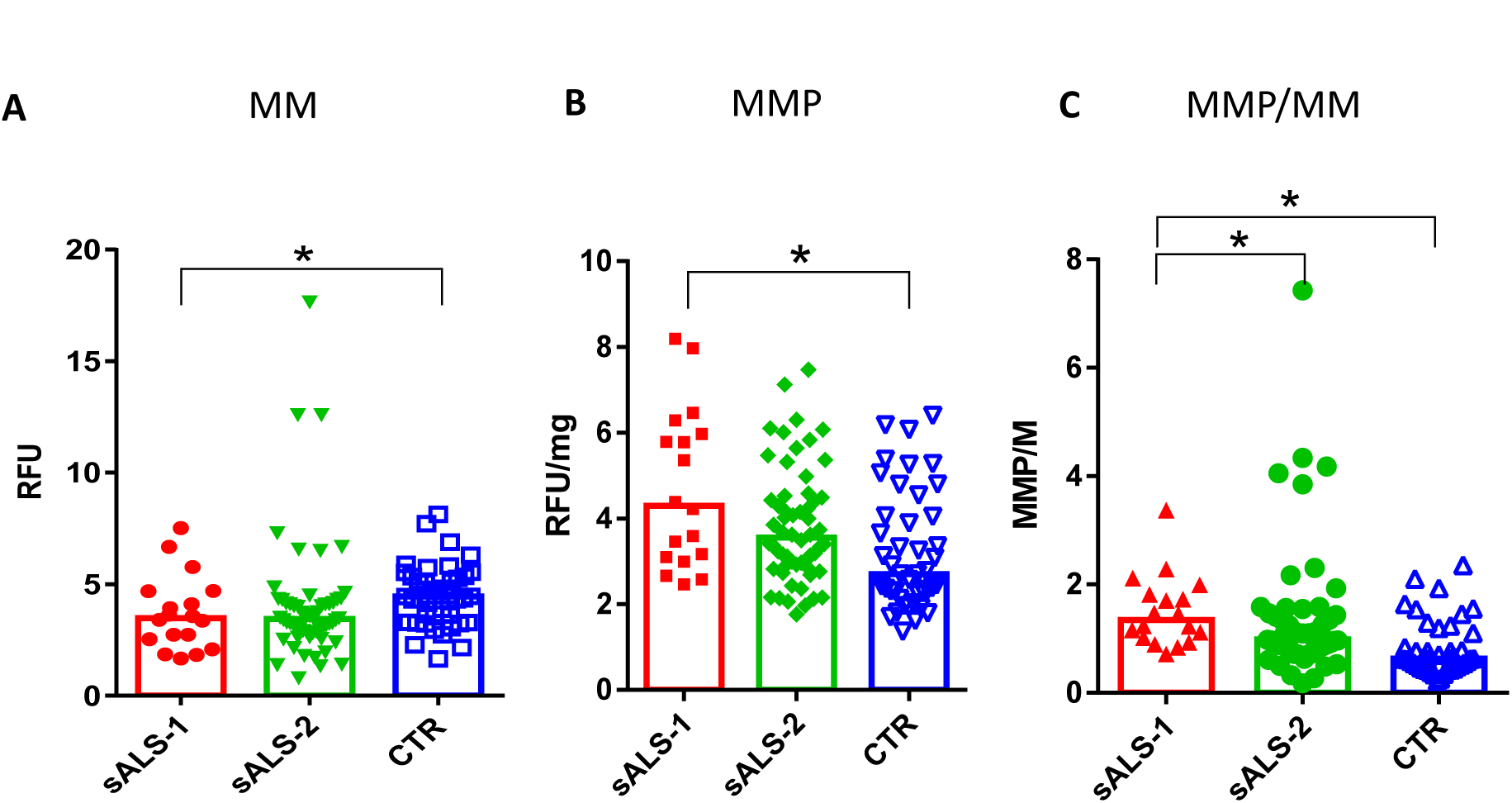
Mitochondria membrane potential in increased in sALS-1 compared to sALS-2 and control. *, P<0.05 (Mann Whitney test), sALS-1, n=18, sALS-2, n=55; controls, n= 43.

**Figure S7.**
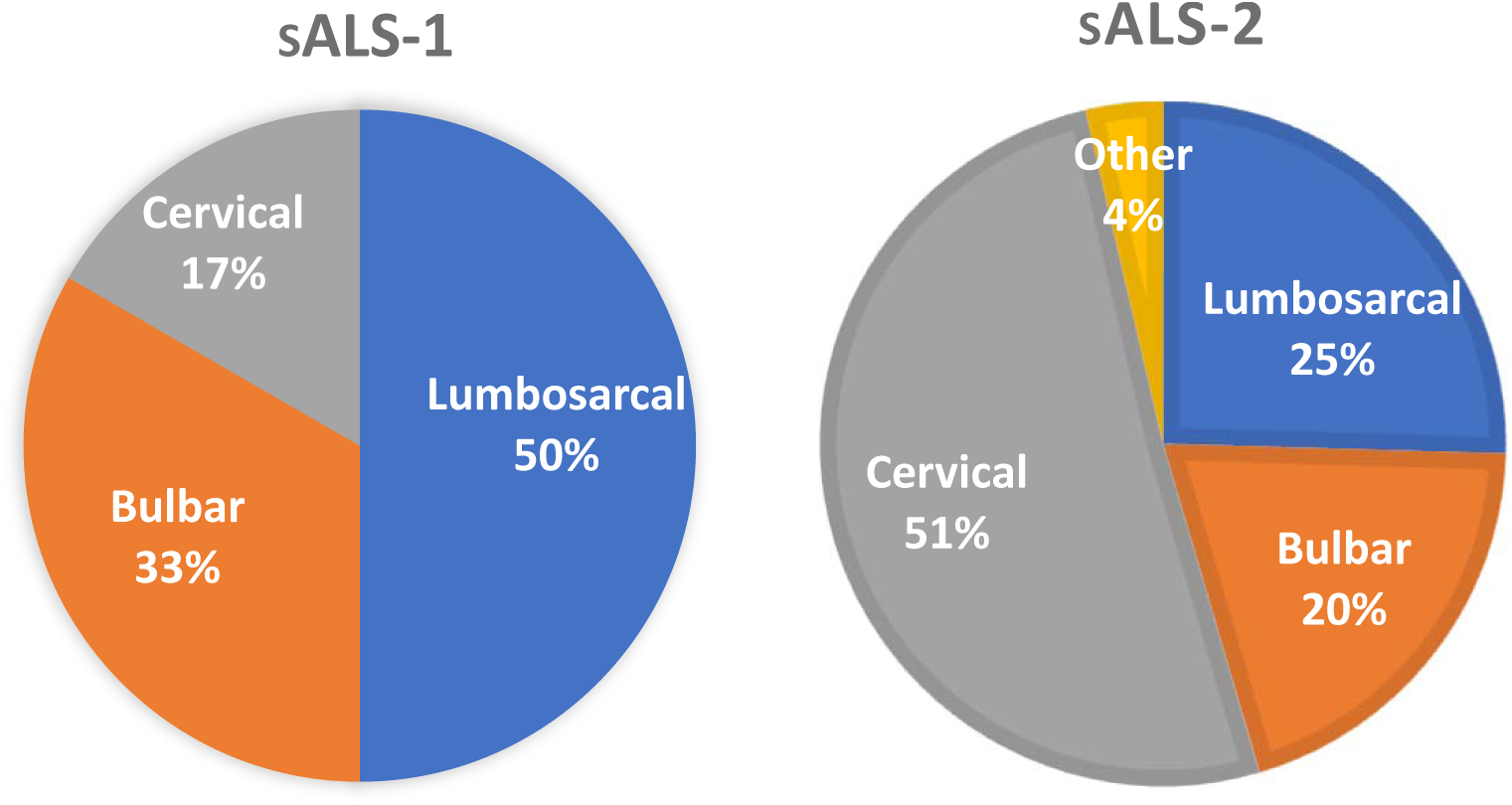
sALS-1 patients show a different frequency distribution in the site of disease on-set compared to sALS-2. sALS-1, n=18; sALS-2, n=55.

